# Polar-angle representation of saccadic eye movements in human superior colliculus

**DOI:** 10.1101/169003

**Authors:** Ricky R Savjani, Elizabeth Halfen, Jung Hwan Kim, David Ress

## Abstract

The superior colliculus (SC) is a layered midbrain structure involved in directing eye movements and coordinating visual attention. Electrical stimulation and neuronal recordings in the intermediate layers of monkey SC have shown a retinotopic organization for the mediation of saccadic eye-movements. However, in human SC the topography of saccades is unknown. Here, a novel experimental paradigm and highresolution (1.2-mm) functional magnetic resonance imaging methods were used to measure activity evoked by saccadic eye movements within SC. Results provide three critical observations about the topography of the human SC: (1) saccades along the superior-inferior visual axis are mapped across the medial-lateral anatomy of the SC; (2) the saccadic eye-movement representation is in register with the retinotopic organization of visual stimulation; and (3) activity evoked by saccades occurs deeper within SC than that evoked by visual stimulation. These approaches lay the foundation for studying the organization of human subcortical eye-movement mechanisms.

**Highlights:**

- High-resolution functional MRI enabled imaging from intermediate layers of human SC
- Saccades along superior-inferior visual field are mapped across medial-lateral SC
- Saccadic eye movement maps lie deeper in SC and are in alignment with retinotopy

**eTOC Blurb:** Savjani et al. found the polar angle representation of saccadic eye movements in human SC. The topography is similar in monkey SC, is in register with the retinotopic organization evoked by visual stimulation, but lies within deeper layers. These methods enable investigation of human subcortical eye-movement organization and visual function.

## Introduction

The superior colliculus (SC) is a layered midbrain nucleus with multiple functions including processing visual input, orienting attention, generating saccadic eye movements and integrating multiple sensory inputs. The SC is organized into seven layers of distinct cytoarchitecture characterized by alternating cellular and fiber layers, with three outer layers comprising the superficial layers and four inner layers (Wurtz and Albano, 1980). The three superficial layers receive direct retinal input and contain a retinotopically organized topography of visual receptive field neurons (Cynader and Berman, 1972; Feldon and Kruger, 1970). The inner layers are separated into intermediate and deep layers. The two intermediate layers contain motor output neurons to control eye movements (Robinson, 1972), whereas the two deep layers contain multisensory integration neurons (Meredith and Stein, 1990; Sprague and Meikle, 1965).

The oculomotor function of the SC was first discovered by electrical stimulation (Adamuk, 1872; Donders, 1872), and later induced via strychnine (Apter, 1946) in anesthetized cats. Extensive studies in alert monkeys via electrical stimulation (Robinson, 1972), neuronal recordings (Mohler and Wurtz, 1976), and lesions (Wurtz and Goldberg, 1972) then characterized the non-human primate topography of the SC [see several review papers including: (Fuchs et al., 1985; Sparks and Hartwich-Young, 1989; Sparks and Jay, 1986; Wurtz and Albano, 1980)]. In particular, the intermediate layers of monkey SC contain a retinotopically organized saccadic eye-movement map (Schiller and Stryker, 1972). Saccade magnitude is roughly mapped along the rostral-caudal axis of SC, while saccade direction is mapped along the medial-lateral axis of the SC in monkeys. The mapping is not linear, with over-representation of small-magnitude saccades, and of polar angles close to the horizontal meridian (Sparks et al., 1976). Recent studies have further revealed that the upper visual field is over-represented compared to the lower visual field with smaller and more sensitive receptive fields (Hafed and Chen, 2016). Also, the SC topography shows a small but variable amount of tilt in the left-right direction that is in rough correspondence to the anatomic tilt of the colliculi relative to the neuraxis (Robinson, 1972). The output motor neurons of the intermediate layers of the SC project to two distinct nuclei to control vertical eye movements (Horn and Büttner-Ennever, 1998): the rostral interstitial nucleus of the medial longitudinal fascicle that drives saccade initiation (Büttner et al., 1977) and the interstitial nucleus of Cajal, which is involved in integrating eye vertical velocity and position (Crawford et al., 1991).

In awake, behaving humans, studies to infer SC function have been largely limited to lesion analysis and whole-brain functional magnetic resonance imaging (fMRI). Lesion studies have confirmed some functions of human SC as compared to animal models and continue to provide valuable information (Sereno et al., 2006; Biotti et al., 2016). However, it is rare to find human patients with focal lesions to the SC without comorbid complications. Further, some aspects of saccadic eye movements can recover even after direct lesions to monkey SC (Hanes et al., 2005). Whole-brain or low-resolution (i.e., ≥2-mm voxels) fMRI has documented saccade-related activity in SC (Furlan et al., 2015; Krebs et al., 2010; de Weijer et al., 2010), and reach-related functions performed by the deep layers of SC (Himmelbach et al., 2007; Linzenbold et al., 2011; Linzenbold and Himmelbach, 2012; Himmelbach et al., 2013). However, the low-resolution measurements could not delineate the detailed topography of SC functions.

Previous human studies of saccadic mapping in cortex have attempted to use fMRI phase-encoding approaches (Connolly et al., 2015; Konen and Kastner, 2008; Schluppeck et al., 2005; Sereno et al., 2001); however, these designs had two critical limitations for imaging subcortical activity: (1) a very low duty cycle (1 saccade every 5 s) and (2) reverse saccades made immediately after forward saccades. The low duty cycle forces participants to fixate most of the time instead of making saccades, which reduces the evoked hemodynamic activity (**Figure S1A**). Further, if participants are instructed to make a saccade in the opposite direction of the target, these reverse saccades may resemble an anti-saccade, or an eye-movement away from the cued target. Recent fMRI studies in humans revealed that collicular activity elicited from a saccade towards a target (a pro-saccade) was not distinguishable, via pattern recognition, from anti-saccades (Furlan et al., 2016), suggesting that such anti-saccades could alter topographic measurements. Also, performing anti-saccades may involve the release of inhibitory control exerted by frontal regions like the dorsal lateral prefrontal cortex (DLPFC) onto the colliculus (Condy et al., 2004), which may confound SC topography measurements (**Figure S1**). A paradigm with a higher duty cycle that isolates saccades in one direction is crucial to getting reliable SC topography measurements with fMRI.

Recent work was able to demonstrate the presence of visual stimulation retinotopy (Schneider and Kastner, 2005), and covert attention signals (Schneider and Kastner, 2009) in the superficial layers of SC using higher resolution fMRI (1.5 x 1.5 x 2mm voxels) targeted specifically to midbrain. Previously, our laboratory has used still higher resolution fMRI (1.2-mm cubic voxels) to measure how visual attention and stimulation are arranged on the superficial and intermediate layers of SC (Katyal and Ress, 2014; Katyal et al., 2010, 2012), confirming macaque electrophysiology measurements and the registration of visual stimulation and attention. Here, we improved those methods and developed a novel task to measure the polar angle representation of saccadic eye movements, elucidating their topography within the intermediate layers of human SC.

## Results

### Eye-movement paradigm design

We designed a paradigm in which participants performed many saccades in one direction while minimizing saccades in the opposing direction (see **Movie M1**). Participants made saccades either to the left or to the right (activating primarily the contralateral SC) while we cyclically varied the vertical component of the saccade to correspond to the lower, horizontal, and upper visual field (**Figure 1**). Participants performed three 6° saccades guided by a green dot target in a static grid of 12 red dots. The static red dots were arranged with 4 dots separated by 6° along each of the three principle axes (horizontal, 45° diagonal, and −45° diagonal). The use of a static grid reduced differential contrast effects from retinal slip. The use of green-red color contrast minimized the effects of bottom-up color contrast in target discrimination. Further, human SC has recently been shown to adapt to red-green contrast (Chang et al., 2016), so our static red-green grid reduces cue-evoked visual stimulation during saccadic eye movement measurements.

**Figure 1.**
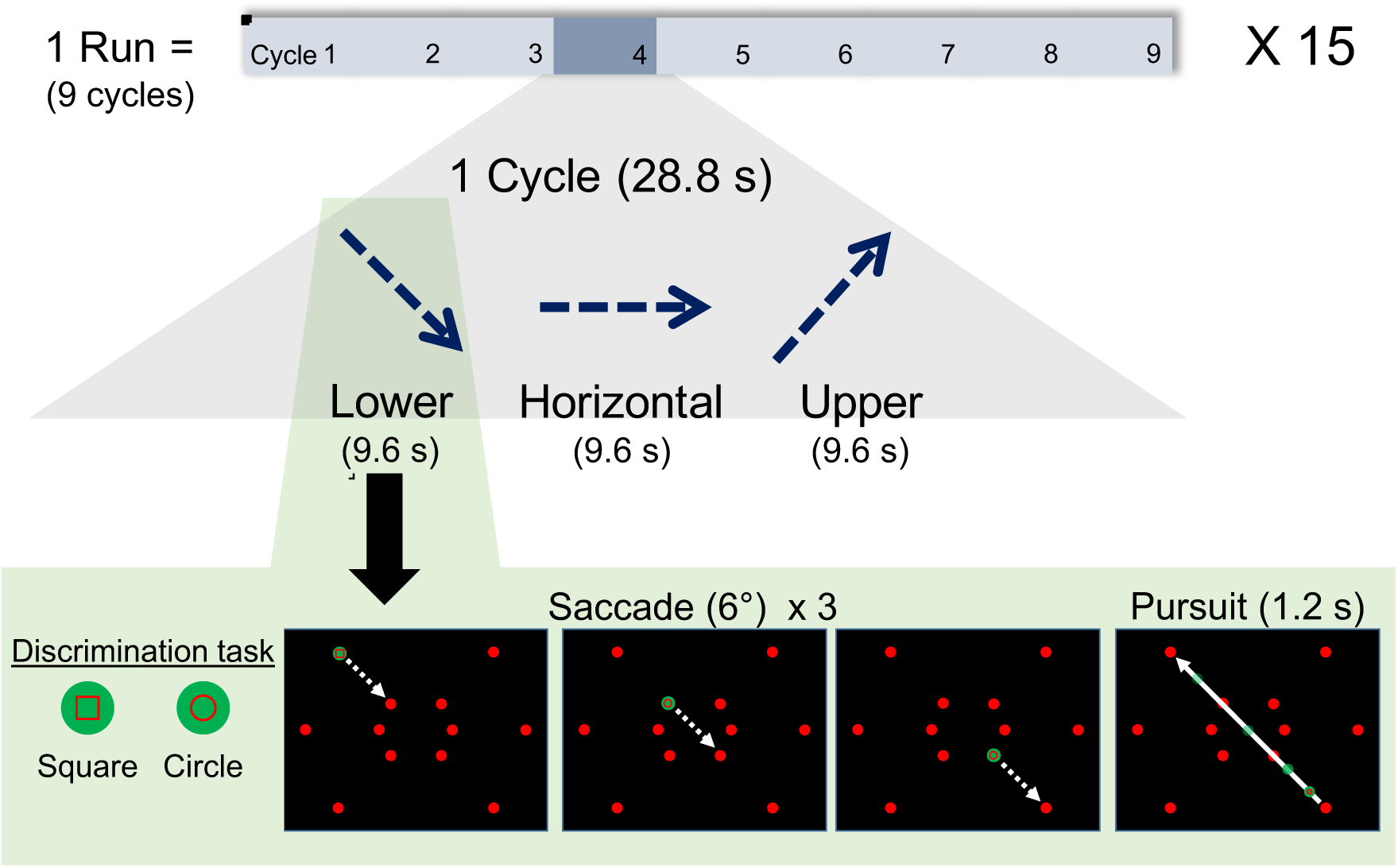
Participants performed visually-guided saccades to measure the polar-angle representation of saccades in SC. In each session, activity from one SC was measured by having participants perform saccades toward a single hemifield (right shown here) along a particular polar angle. The stimulus screen showed a static grid of 12 red dots with one target dot turned green to indicate the saccade target. Participants made three 6° saccades along the current polar angle, after which a 1.2-s visually-guided smooth pursuit was made back to the origin along that axis. Upon fixation onto target dots and during the smooth pursuit, participants performed an object discrimination task (square or circle) to keep attention engaged and improve the reliability of eye movements. In each 28.8-s cycle, the vertical component of the saccades progressed through three polar angles: –45° (lower), 0° (horizontal), and +45° (upper). Each session consisted of 15 ~4.5-min runs, each of which included 9 cycles. (See **Figure S1** for evolution of task design).

Participants initially held fixation at an upper corner of the display. The first saccade was initiated when an adjacent red dot turned green to indicate the saccade target. Once the saccade was made, participants had to discriminate between the outlines of two possible shapes (circle or square) presented within the target dot. This required visual attention to be engaged and saccades to be made more reliably. Participants responded via button press, which triggered the green dot to appear at the next target along the current axis. After three saccades, the participants then performed a smooth pursuit (1.2 s) back to the first dot. Saccades and the pursuit were continued along the same axis for 9.6 s, and then participants performed another smooth pursuit to the start of the next axis. During each smooth pursuit, the discrimination task had to be performed 0—2 random times (truncated Poisson distribution, λ = 1) during the 1.2-s smooth pursuit, encouraging attention to remain engaged and eye movements to be restrained to the pursuit path. Participants performed 9 cycles in a single run (~4.5 min) and ~15 runs per session. Leftward and rightward saccades were run on separate sessions to measure the contralateral response of each SC independently.

### Behavioral performance

During the eye movement task, participants performed the object discrimination task reliably (mean 82.8 ± 8.6% across all sessions and participants, **Figure 2A**). The frequency of the saccades was determined by the participants, as the button response in the discrimination task triggered the onset of the next saccade cue. Participants generally made saccades with an interval between 0.6 – 0.8 s (**Figure 2B**). During the visual stimulation protocol, participants performed the speed discrimination task at a mean accuracy of 70.3 ± 5.1% (**Figure 2C**), close to the expected staircase asymptotic accuracy of 71%, with a mean discrimination threshold of 1.6 °/s (**Figure 2D**).

**Figure 2.**
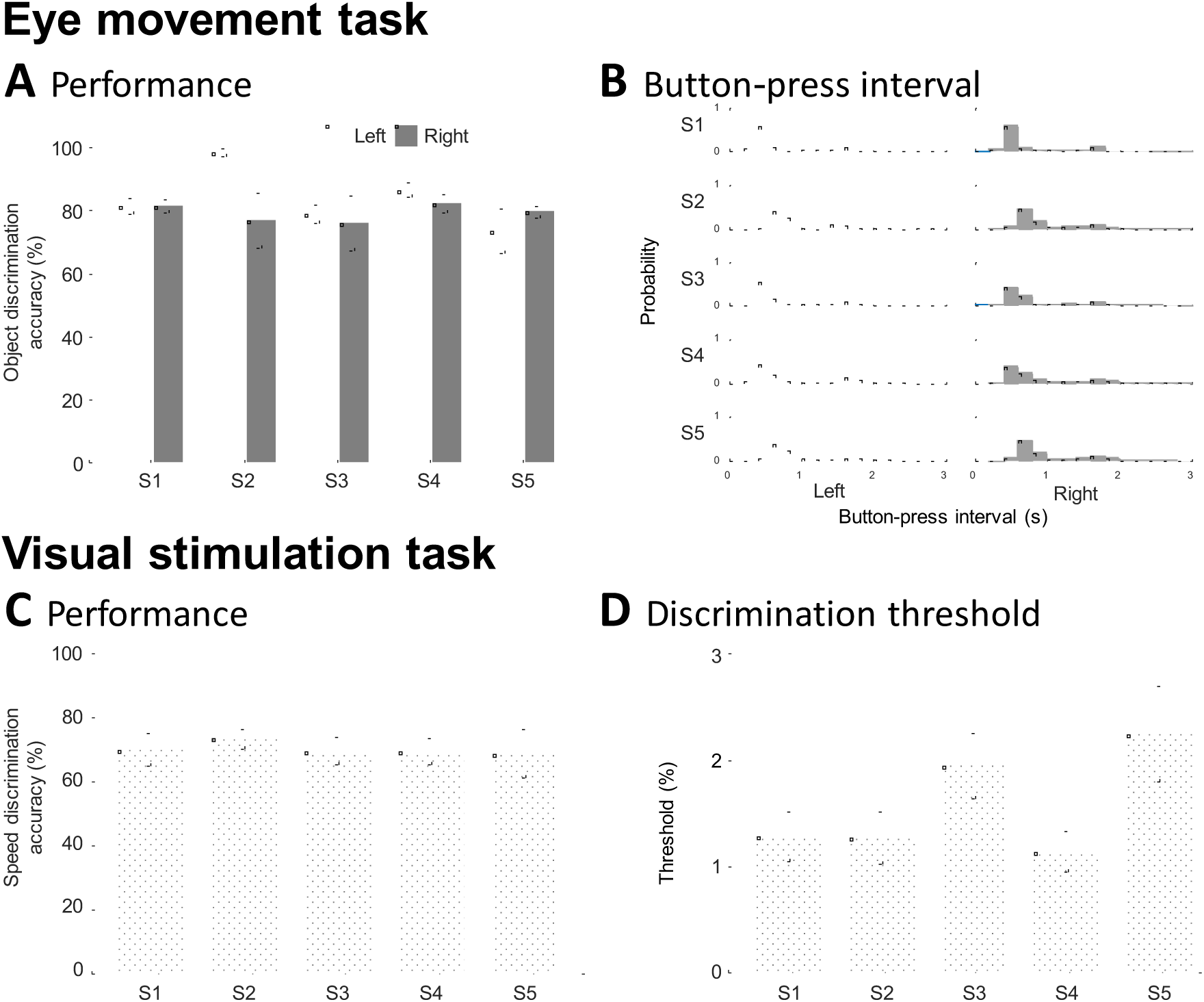
Psychophysical measures show that participants reliably performed both the object discrimination task during saccadic eye movements (top row) and the speed discrimination task during visual stimulation (bottom row). **A.** Participants detected the object (circle or square) with 82% accuracy. Histograms of the button-press intervals are bi-modal, with one mode around 0.6 – 0.8 s corresponding to the saccades and a slower mode at 1.2 – 1.5 s corresponding to the smooth pursuit. (**B**) Participants performed the speed-discrimination task at ~71% accuracy and mean discrimination threshold of 1.6 °/s. Error bars show standard deviation across all trials.

### Eye movements

Before each saccade-mapping session, we trained all participants on 2—3 runs outside of the scanner and quantified the reliability of their eye movements (**Figure 3A**). Saccades were detected and visualized on polar plots to show the eccentricity and polar angle of each saccade. Saccades were color coded to represent their cycle timing along the three polar angles of the task. We were also able to obtain reliable eye tracking in the scanner from two participants on both rightward and leftward eye-movement sessions (**Figure 3B**). Histograms confirm eye movements slightly hypermetric to the cued amplitude (6.29±1.95°, shown in gray) in the cued direction (angular error: 0.82°± 10.01°). Longtailed distributions in the opposite direction (white) were also observed. The task periods designed to evoke smooth pursuits often contained saccades of variable amplitude, but most were <3° (median: 0.8°, mean: 3.07°±3.19°). Small correction saccades also contribute to the opposite-direction saccades, as saccades to targets were often hypermetric, followed by a small correction saccade.

**Figure 3.**
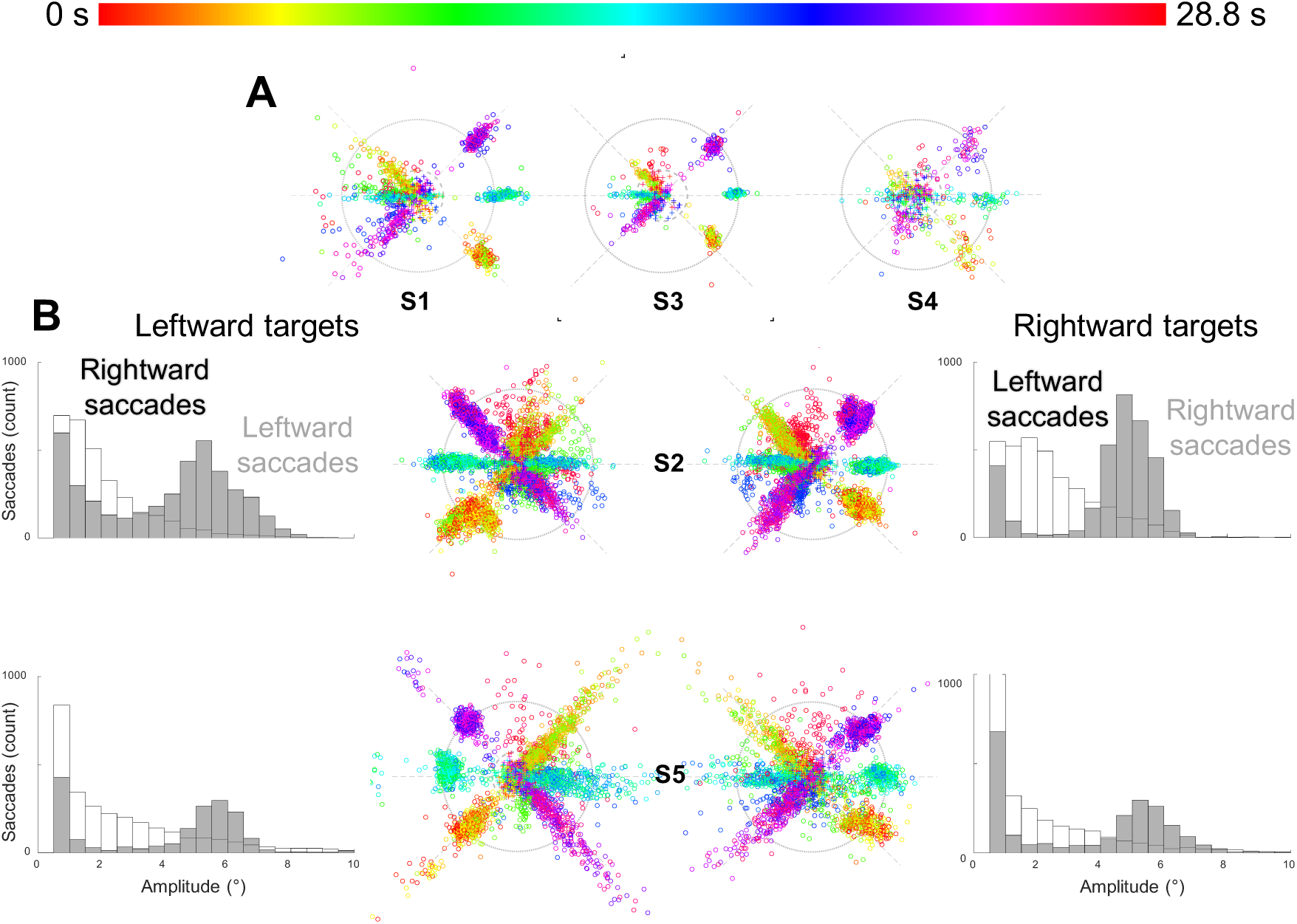
Eye tracking confirmed the reliability of participant’s eye movements. **A.** Data from three participants show saccades during a training session outside of the scanner. Polar plots show the polar angle and eccentricity of each saccade detected, as well as the corresponding time during the cycle, represented as a color from the HSV color map. **B.** Data from two participants during fMRI acquisition inside the scanner also show reliable eye movements during both leftward and rightward sessions. Histograms show eye movement amplitudes of 6.29° ± 1.95° in the cued direction (shown in gray). Long tailed distributions in the opposite direction (white) were also observed (median: 0.8°, mean: 3.07° ± 3.19°).

### Surface Analysis

Using the 0.7-mm voxel structural volume, we segmented the brainstem (including portions of the thalamus) using a combination of automatic (e.g., active contour evolution) and manual approaches in ITK-SNAP (Yushkevich et al., 2006) (**Figure 4A**). Two participant’s brains were initially segmented using newer Bayesian approaches in FreeSurfer 6.0 (Iglesias et al., 2015), followed by manual editing. A surface model was built at the tissue-cerebrospinal fluid interface using a deformable surface algorithm (Xu et al., 2006). Functional data were then spatially aligned and resampled to the high-resolution T1 volume, averaged across runs, and visualized on the surface. Distance along the surface, *d_mesh_*, was estimated by summing Euclidean distances along the mesh from a reference point on each participant’s inter-collicular axis. A Euclidean nearest-neighbor distance map was also computed from SC tissue voxels to the vertices of the surface to give a measure of the depth (*s*, mm) of the tissue voxels (Khan et al., 2011; Ress et al., 2007). Time-series data were generally averaged over a particular range of depth values to improve SNR.

**Figure 4.**
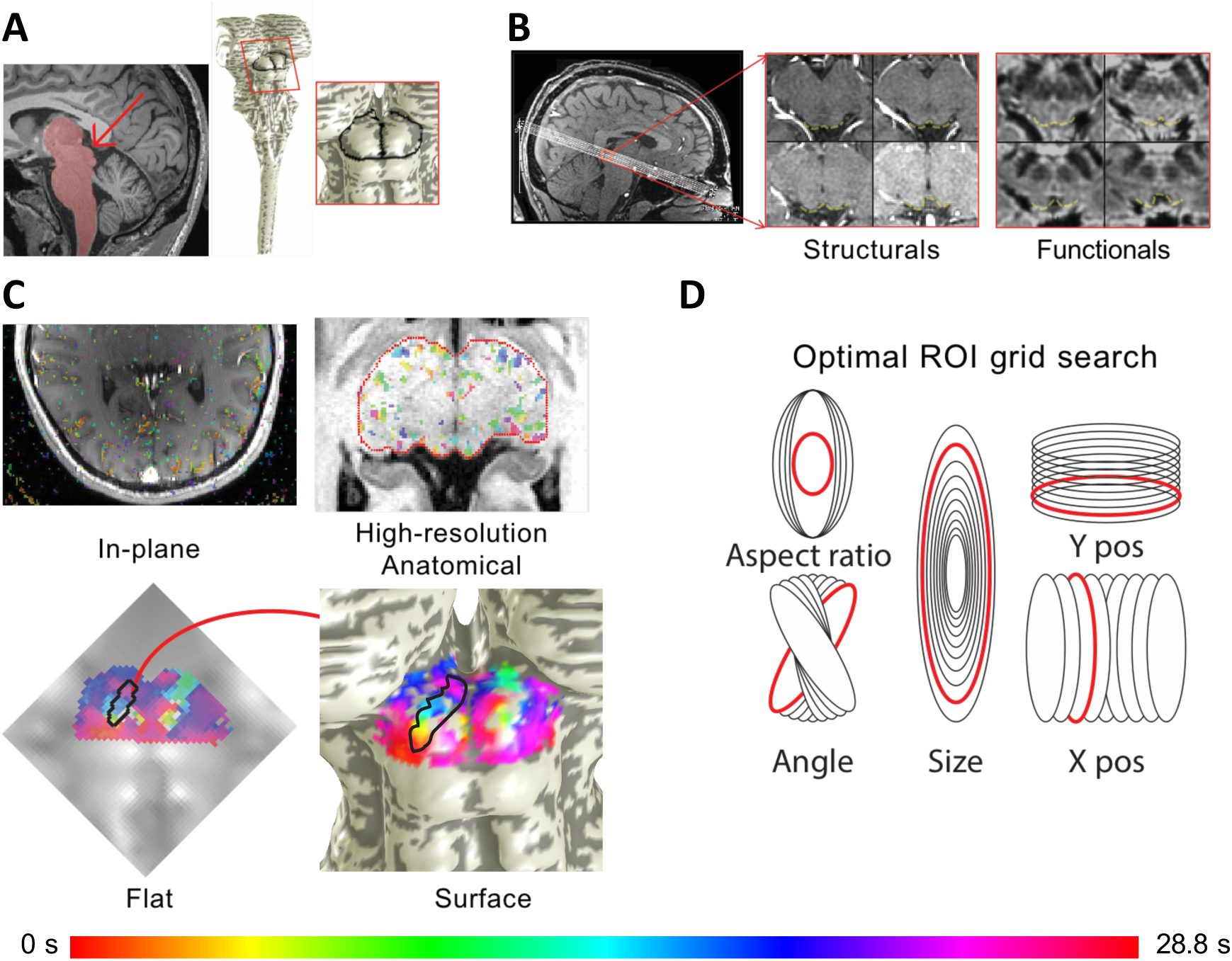
SC image acquisition, segmentation, surface projection, and flat maps allowed data visualization to elucidate phase progressions, and optimal selection of ROIs delineating saccadic eye movement representations. **A**. High-resolution (0.7 mm) T1 anatomical images were used to segment out the brainstem and create surface models for each participant. The surface maps allowed visualization of the posterior view of the midbrain and SC. **B**. Eight quasi-axial slices through the SC were acquired. In-plane T1 structural slices with the same slice prescription as functional data were obtained before and after collection of functional runs to facilitate registration to high-resolution T1 anatomical. **C**. Functional data were phase-mapped and the best fitting phases for each voxel were overlaid on the in-planes and then registered and overlaid onto the high-resolution anatomical volume, which could then be projected onto the surface by depth averaging. The depth-averaged functional data were also visualized and analyzed on a 2D projection. **D**. In the flattened view, thousands of elliptical ROIs were generated by manipulating 5 elliptical parameters. The ellipse that best fit the optimization criteria was visualized in the flat view to check for accuracy and then transformed into the volume and the surface model.

### Phase Mapping

A sinusoid at the stimulus repetition frequency (24 s for visual stimulation data, 28.8 s for eye movement data) was fit to the depth-averaged data (0-1.6 mm for visual stimulation data, 0.6-1.8 mm for eye-movement data) using coherence analysis. These fits provide three parameters: amplitude, phase, and coherence. Amplitude measures the strength of the evoked activation, phase measures the time delay of the response, and coherence is equivalent to the Pearson’s correlation coefficient, quantifying the fraction of variance explained by the sinusoidal fit. The phase, specifically, is used to measure the relationship of the fMRI response to the current state of the stimulus (e.g., saccade angle, or visual stimulus polar angle). Phase maps, expressed in units of time delay, were projected onto the brainstem surface to visualize the topography of visual stimulation and eye movements.

### ROI generation

We generated elliptical ROIs to demarcate the phasic progressions of saccade-evoked activity. First, surface representations of the SC were flattened to form a 2D image (Wade et al., 2002; Wandell et al., 2000) (**Figure 4C**). The phasic progression from medial to lateral was visually observed and then delineated with two vertices on the surface to define the start and stop of the putative eye movement maps; these two vertices were then transformed to the flat image. Next, many (~20,000) elliptical ROIs were generated by varying 5 parameters (**Figure 4D**): size (15 – 60% of one whole SC area); aspect ratio (3.5—7); *x*,*y* center coordinates (each varied ±2 pixels from the midline of the delineated phase progression); and the elliptical angle (±15° relative to the angle of the delineated phase progression). An exhaustive search was then conducted to maximize five criteria: (1) number of activated (*p* < 0.2) voxels, *v_sig_*; (2) geometric average *p*-value, *p̄_geom_* = 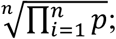 (3) phase range (defined by median absolute deviation), *ϕ_range_* = median(*ph_i_* - median(*ph*)); (4) correlation (variance explained) of the linear fit between the mesh distance *d_mesh_* and the phasic progression 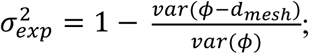 and (5) the reciprocal deviation of the same linear fit and the expected progression across the collicular width, *δ* = 1/*abs*(*ellipse_slope_* – *target_slope_*). The five optimization criteria were multiplied together to yield an overall metric, *opt* = 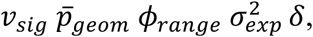 for each ROI. The top four values of *opt* identified the best ROIs, which were examined visually for quality confirmation. The top fitting ROI was used for subsequent analyses.

### Polar angle maps of saccadic eye movements

Saccades along the superior-inferior visual field were mapped along the medial-lateral axis of the SC in all 5 participants (**Figure 5**). Our eye-movement task involved making 6° saccades toward one hemifield, with smooth pursuit in the opposite direction.

**Figure 5.**
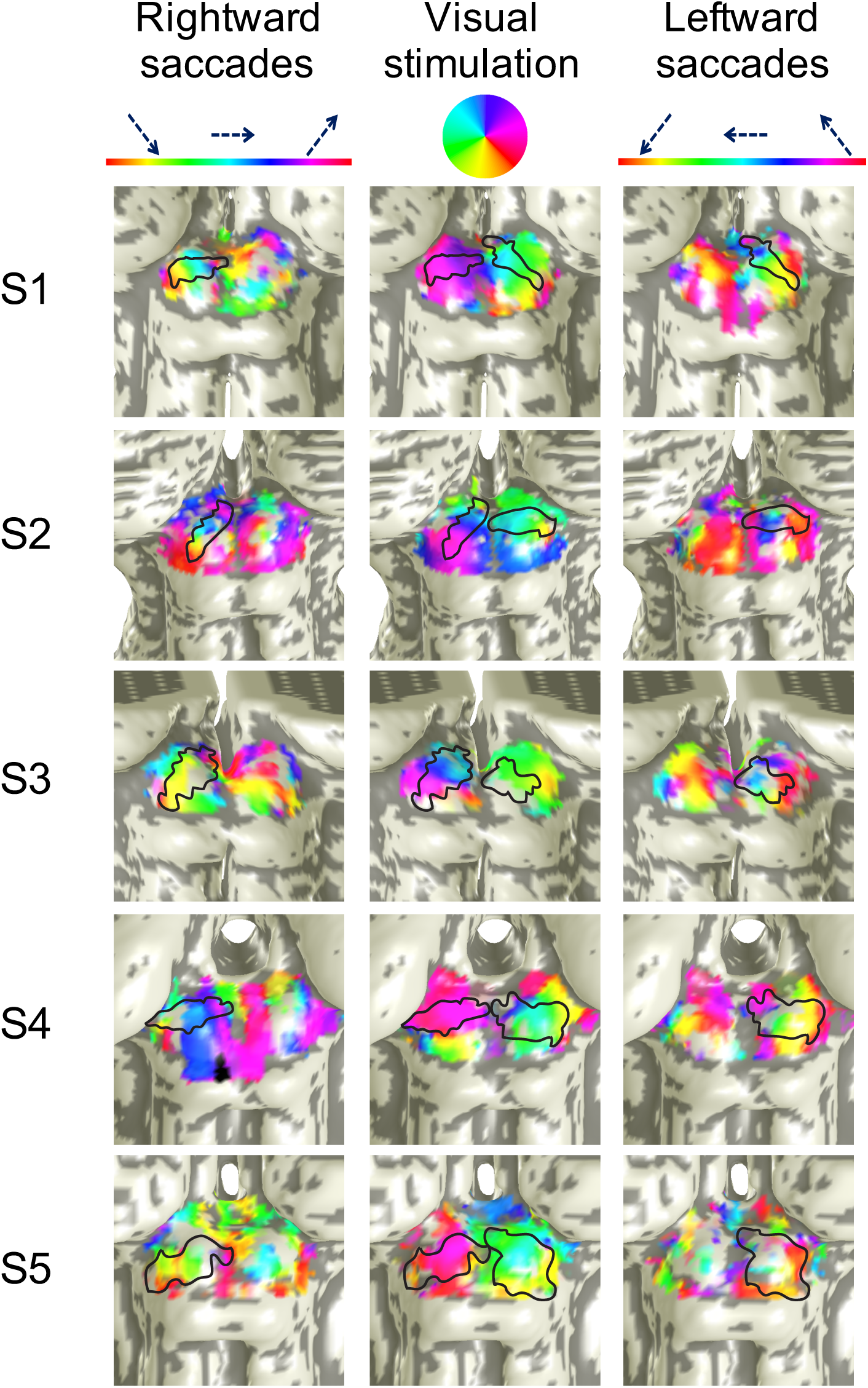
Saccadic eye movements along the superior-inferior visual field were mapped medial-laterally on the contralateral SC in five participants. From left to right, the columns of SC surfaces show phase progressions of rightward saccades (left SC), retinotopic mappings, and leftward saccades (right SC). Regions outlined in black show the optimally defined elliptical ROIs on each SC. The elliptical ROIs are also shown on the retinotopy for comparison. (See **Figure S3** for cortical analyses).

Accordingly, we see organized phase progressions only in one SC at a time. ROIs, obtained by the aforementioned optimization procedures, capture the range of these phase progression evident near the crown of the colliculus. This is quantified by significant correlations of saccadic eye-movement phase with distance along the medial-lateral axis of the SC at the group level (slope: 0.451, *r* = 0.734, *p* ~ 0.0002, **Figure 6**). Significant correlations (*p* < 0.05) were also observed in 7 out of 10 individual SCs (**Figure S2**). Strong activity was also evident in many other portions of both contralateral and ipsilateral colliculi. In particular, several ipsilateral colliculi exhibit reversed phase progressions (S1L, S2L, S4L, S1R, S2R, S3R, and S4R) that were more rostral than the contralateral progressions. This rostral pole activation could represent activity from the smooth pursuits, but in our experience, the rostral pole is often contaminated with vascular nuisance activations, so these activations will not be further discussed.

**Figure 6.**
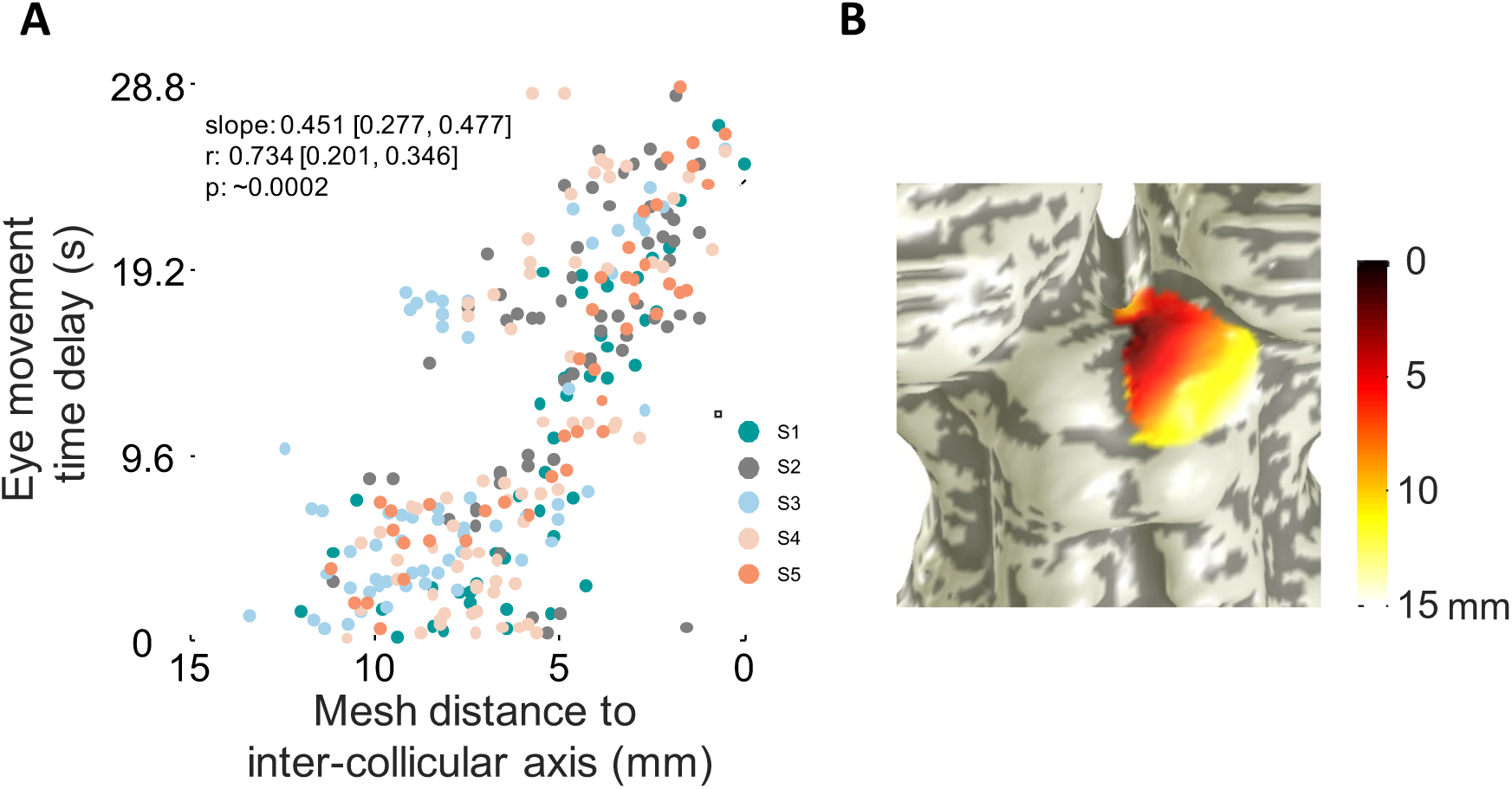
Saccadic eye-movement phase increases medially, revealing saccades made superiorly are mapped medially. **A.** The phase of all active vertices (*p* < 0.2) is plotted against mesh distance from inter-collicular axis for all participants combined, revealing the lateral-medial mapping of eye-movements along the inferior-superior visual axis. Correlation metrics are listed, with 68% confidence interval ranges of slope and correlation values listed, as well as bootstrapped p-values. **B.** Representative participant illustrates mesh distance on a surface SC (data from participant S2). (See **Figure S2** for individual participant plots).

Weak but significant activation was also observed in early visual cortical areas V1—3 (**Figure S3**). However, this activity had a biphasic rather than triphasic character. These results suggest a small amount of activation from unbalanced visual motion between horizontal and off-horizontal eye movement phases (see Discussion).

Visual stimulation (with an attentional task) produced clear, largely medial-lateral phase progressions corresponding to a representation of polar angle in the superficial layers of SC (**Figure 5**, center column). The stimulation-evoked phase progressions are qualitatively consistent with the progressions evoked by the saccade tasks in each SC. In most participants, we also observed a small rostral tilt in the lateral-to-medial phase progression. This tilt also corresponded to the tilt in the saccade data. No consistent phase progressions were elicited in early visual cortex (**Figure S2**), indicating that the phase progressions were not associated with visual stimulation alone.

### Correlations between eye-movement and visual stimulation topographies

Within the elliptical ROIs for each participant, we observed the saccadic eye movement phases of the three angles of movement to be in alignment with the visual stimulation retinotopy for both left (slope: 0.733, *r* = 0.312, *p* = 0.0052) and right (slope: 0.928, *r* = 0.567, *p* = 0.0034) SCs (**Figure 7**). Quantitative correlations were significant for five out of ten SCs in individual participants (**Figure S4**).

**Figure 7.**
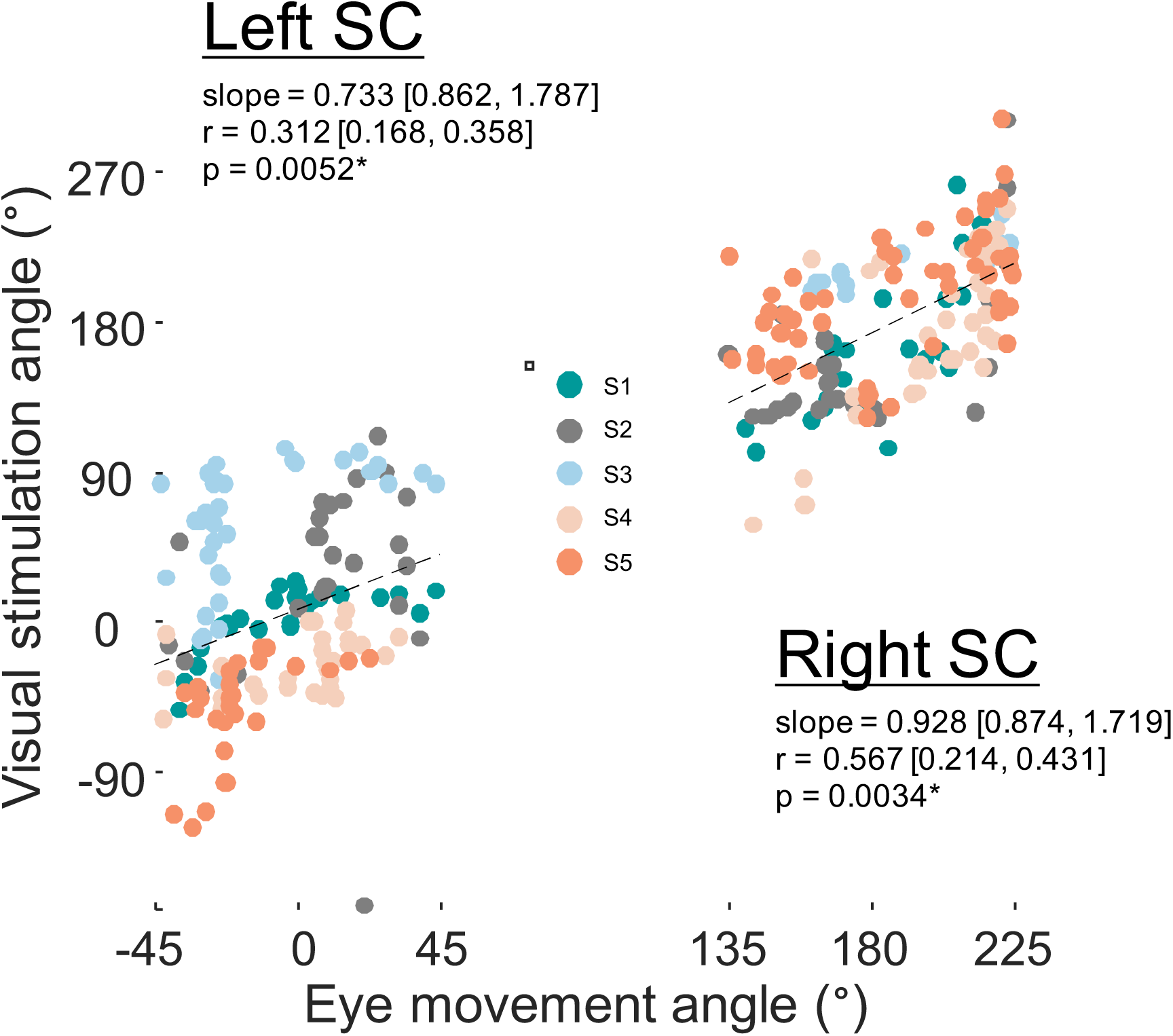
Retinotopic organization of superficial layers of SC corresponded with phase progressions of the deeper saccadic eye movement maps. Significant correlations were found for both left and right SCs collapsed across all participants, indicating that these two maps are in alignment. Correlation metrics are listed, with 68% confidence interval ranges of slope and correlation values listed, as well as bootstrapped p-values. (See **Figure S4** for individual participant’s correlations).

### Depth profiles

Laminar depth profiles extended deeper into SC for activity evoked by saccadic eye movements compared to visual stimulation (**Figure 8**). The peaks are significantly shifted deeper for eye movement maps compared to visual stimulation maps for both left (shift Δ = 0.9 mm, *p* = 0.0012) and right (Δ = 0.7 mm, *p* = 0.0002) SC. At the individual level, data were significant in four out of 10 SC (**Figure S5**).

**Figure 8.**
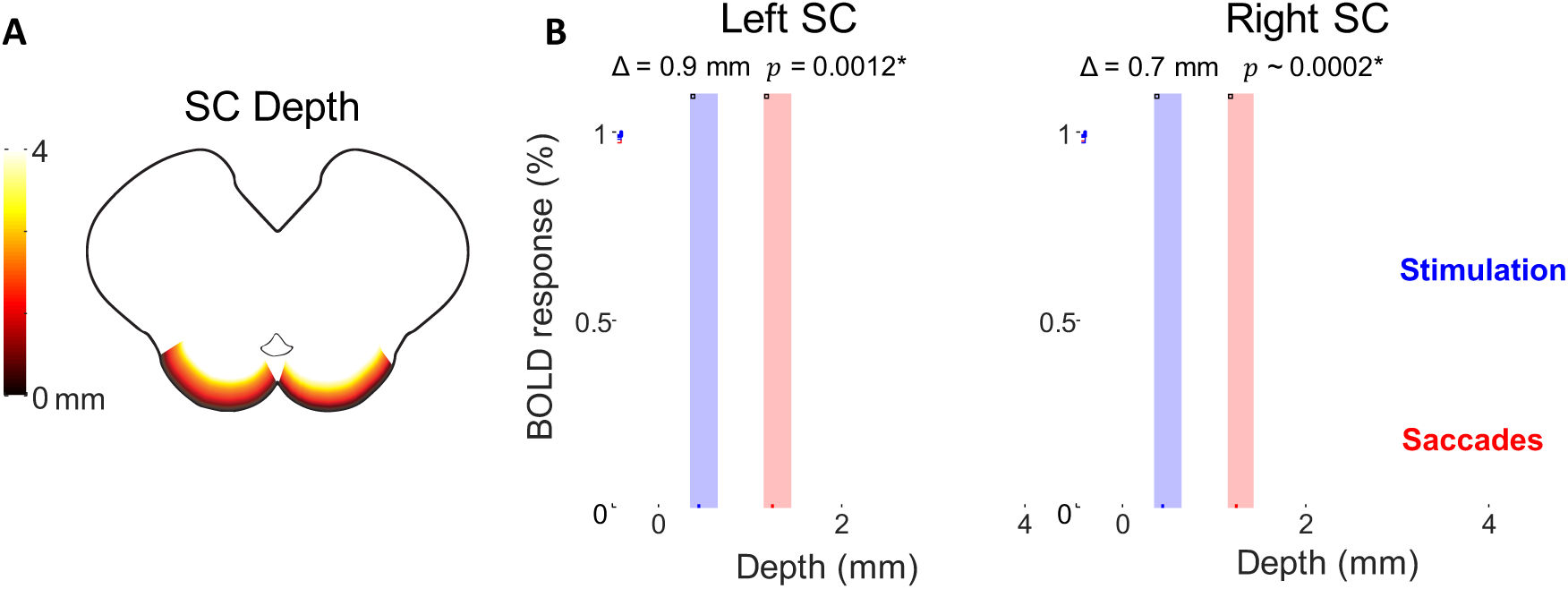
Saccadic eye movement maps lie deeper in SC than visual stimulation maps. **A.** Graphic illustrating depth of SC in a horizontal cross section. **B.** Laminar depth profiles for both left and right SC show the activity evoked by saccades (red) lies deeper in SC than activity evoked by visual stimulation (blue). Data are combined across all 5 participants for left and right SC. Dotted lines represent 68% confidence intervals bootstrapped across all runs and participants. The peak is significantly shifted deeper for saccades for both left and right SC (shaded rectangles show bootstrapped 68% confidence intervals for centroid calculations). (See **Figure S5** for individual participant depth profiles).

## Discussion

We used high-resolution functional MRI to map the polar angle representation of saccadic eye movements within human SC. We found saccades along the superior-inferior visual axis were mapped roughly medial-to-lateral on the anatomy of the SC. Eliciting these maps required a novel paradigm in which participants were cued to make saccades along one of three specified direction axes, and then a smooth pursuit back to the start of the axis. Doing so isolated forward saccades from reverse saccades, while still allowing participants to make many saccades along a controlled set of directions in an experimental session. In addition, we utilized a fixed and symmetric grid of saccade targets to minimize the effects of motion and contrast evoked by retinal slip. This experimental design was not immediately obvious to us but evolved over time and was only discovered after many failed paradigms (**Figure S1**).

Our experimental stimulus still evoked some activity beyond the contralateral medial-lateral phase progression. In 7 out of 10 sessions, weaker but significant reverse phase progressions appeared on the ipsilateral SC. There are at least three possible reasons for this ipsilaterally evoked activity. First, this activity could be driven by small saccades during the smooth pursuits, particularly as the pursuit approaches the target endpoint, and the participant makes a predictive saccade ahead of the pursuit. Second, the ipsilateral activity may be evoked by the observed small correction saccades, as the participant often saccades past the target and corrects by performing one-or-more saccades in the opposite direction to reach fixation upon the target Third, it may be that the prior targets are still remembered during the smooth pursuit; activity in monkey SC of both remembered and visually-guided saccade targets has been reported during such smooth pursuits (Dash et al., 2016)..

We found that the representation of saccades in human SC to be in good alignment with the overlying retinotopic topography, consistent with the topography of monkey SC (Schiller and Stryker, 1972; Wurtz and Goldberg, 1972; Basso and May, 2017). Further, our previous studies showed that superficial attentional maps in human SC are also in alignment with retinotopic maps (Katyal et al., 2010), revealing that attention, retinotopy, and saccadic eye-movement maps all utilize a similar topography.

The depth profiles show that the saccadic eye movement maps lie 0.7—0.9 mm deeper than visual stimulation maps on both SC. This result is consistent with laminar organization inferred from human SC cytoarchitecture (Qu et al., 2006), from human SC response to visual stimulation (Zhang et al., 2015), and from laminar topography across several species including: guinea pig, cat, galago, macaque, and gray squirrel (May, 2006). Profiles showed significant depth separations for only 4/10 participants individually, but very strong differences were seen on aggregate. This reflects the noisy and blurry character of our hemodynamic correlates of neural function; many sessions were also needed to resolve differences between the laminar profiles of attention and visual stimulation (Katyal et al., 2010).

We utilized a fMRI acquisition sequence that pushed the boundaries of 3T subcortical imaging. Our dual-echo spiral sequence (Singh et al., 2017) gave us the higher CNR needed to resolve the intermediate layers of SC. Each session included nearly 2 hours of scanning while the participant performed 4000—5000 saccades – a demanding but feasible physical effort. Our scanning methodology should allow further experimentation of visual-motor and multisensory function of the SC. For example, future work could study the topography of auditory signal representations and multisensory integration in the deeper layers of SC, and test the possibility of ocular segregation in the superficial layers. In addition, our experimental paradigm paves the way for probing topography in cortical frontal eye fields, as well.

There are some limitations to our approach and results. First, saccadic eye movements were not fully controlled. Participants automatically tended to make undesired saccades because of error-correction or prediction. These tendencies broadened the distribution of saccades, and probably generated weaker activation upon the opposite colliculus. Second, visual contrast was not fully balanced as a function of polar angle, resulting in weak activation in early visual areas (**Figure S4**). This weak activation probably reflects the small asymmetry of our fixed cue display. Ideally, this display should have been round, but the display slightly clipped along the top and bottom (**Figure 1**). Therefore, off-horizontal eye movements would create less visual motion drive than on-horizontal movements. This choice was forced by the 16:9 aspect ratio of our MRI-compatible LCD display, and the need to maximize display visual angle to permit sufficient duty cycle. The biphasic activation may have slightly affected our results, but we believe that their effects were comparatively weak given the quality of our observed correlations.

In summary, our techniques allowed us to measure the functional topography of saccadic eye movements on the human SC. This required using high-resolution functional MRI to reliably resolve the laminar nuclei of midbrain, as well as a novel experimental paradigm that allowed participants to make many saccades in one principle direction while minimizing visual stimulation confounds. The topography of the vertical eye movement maps was arranged medial-laterally, in register with the retinotopic representation of visual stimulation, and emerged from deeper layers in SC, similar to the organization observed in monkey.

## Footnotes

**Author Contributions** Conceptualization, R.R.S. and D.R.; Methodology, R.R.S., E.H., and D.R.; Investigation, R.R.S., E.H., J.H.K., and D.R.; Writing – Original Draft, R.R.S. and D.R.; Writing – Review & Editing, R.R.S., E.H., J.H.K, and D.R.; Funding Acquisition, D.R.; Resources, D.R.; Supervision, J.H.K. and D.R.

## Acknowledgements

This work was supported by grants from the NSF (1446377 to D.R.). We would like to thank all members of the CAMRI team at Baylor College of Medicine for housing and facilitating our experiments. Thanks also to Mike Beauchamp and Scott Stephenson for useful feedback on this project.

## Methods

### Participants

We recruited five participants (4 males, all right-handed) to undergo several ~2 hour long scanning sessions. One-to-two sessions were acquired for each participant to discern the polar-angle representation of leftward and rightward saccadic eye movements separately, which were expected to evoke activity primarily in the contralateral SC. Each eye-movement session consisted of 12—16 278-s runs. One-to-two scanning sessions were also acquired from each participant for visual stimulation retinotopic mapping. Visual stimulation experiments evoked activity from both SC in each session. Retinotopy sessions consisted of 14-16 228-s runs. Participants gave informed consent prior to scanning based on our approved protocol from the Baylor College of Medicine Institutional Review Board.

### Stimuli

Stimuli were generated using MATLAB R2015a (Mathworks, Natick, MA) and PyschToolbox-3 (Brainard, 1997) on a Windows 7 Dell PC. Stimuli were presented on a 32” LCD BOLD Screen (Cambridge Research Systems, Kent, UK) at the back of the scanner bore 1.3 m away from the participants’ eyes. The display was gamma corrected using an i1 Pro 2 spectrophotometer (X-Rite, Grand Rapids, MI), and had a mean luminance of 305 cd/m^2^.

### Retinotopy

Retinotopic maps were also acquired for all 5 participants using our previous phase-encoded visual stimulation paradigm (Katyal et al., 2010). Briefly, participants fixated at the center of the display while a wedge (90° polar angle) of moving dots rotated around the entire polar angle to measure retinotopy for both SC in a single session. The rotating wedge consisted of 6 virtual sectors, and in one sector, the dots were moving either faster or slower than all other sectors. Participants performed a speed discrimination task every 2 s. The speed difference was adjusted using a staircase design to control task difficulty and engage attention, boosting signal levels for the retinotopy measurement.

### MRI methods

Imaging was conducted on a Siemens (Erlangen, Germany) 3T Magnetom Trio scanner at the Core for Advanced Magnetic Resonance Imaging (CAMRI) at Baylor College of Medicine. Eight 1.2-mm-thick quasi-axial slices (170-mm field of view) covered the entire SC with the prescription oriented roughly perpendicular to the local neuraxis. Functional data were acquired using a 3-shot spiral (Glover, 1999; Glover and Lai, 1998) dual-echo (both outward) sequence (Li & Glover, 2003) that has been shown to boost functional signal-to-noise ratio (SNR) by >50% compared to single-echo methods (Singh et al., 2017). We used *T_R_* = 0.8 s for each shot, yielding a 2.4-s volume-acquisition time for our 3-shots. To obtain best contrast, *T_E_* was set to 25 ms for the first echo, and acquisition time for each echo was 34 ms, giving an effective echo time of 59 ms for the second echo. The dual echoes were combined as a signal-weighted average.

A set of *T*_1_-weighted structural images was obtained on the same prescription at the end of the session using a three-dimensional (3D) fast low-angle shot (FLASH) sequence (minimum TE and TR, 15° flip angle, 0.9-mm inplane pixel size) (**Figure 2B**). These images were used to align the functional data to the segmented structural reference volume. A high-resolution (0.7-mm cubic voxels) T1-weighted structural volume anatomy was obtained for each participant in a separate session, using an MP-RAGE sequence (TR = 2600 ms, minimum TE, TI = 1100 ms, flip angle = 9°, 3 averages, 24min acquisition time).

### Image data analysis

PREPROCESSING. Image analysis was conducted using the mrVista software package (http://web.stanford.edu/group/vista/cgi-bin/wiki/index.php/MrVista) and a host of custom modifications designed to enable high spatial resolution sub-cortical imaging.

The first 12 s of each retinotopy run were discarded to remove transient effects. Similarly, the first 19.2 s of each eye movement run were discarded to remove transients and to allow participants to become accustomed to the eye-movement task. The following preprocessing was conducted on each of the two echoes for all runs (visual stimulation and eye movement), analogous to our previous pipeline (Katyal et al., 2010): slice-timing correction (zeroed to task onset) and intensity-based motion compensation using a robust algorithm (Nestares and Heeger, 2000). Motion compensation information was obtained only from the first echo because of its higher SNR; these data were then applied to both echoes. The two processed echoes were then combined as a signal-weighted average. Finally, baseline trend removal was performed using a high-pass filtering approach.

LAMINAR PROFILE ANALYSIS. We then examined the amplitude of the complex response as a function of laminar depth within the elliptical SC ROIs, similar to our previous approaches (Katyal et al., 2010, 2012; Katyal and Ress, 2014). Complex amplitude data were first averaged together across all runs for each participant. A phase normalization was performed to correct for the variable hemodynamic time delay across the ROI. In each cylindrical volume element defined by the surface mesh, we obtained the amplitude-weighted mean phase. Complex amplitudes within each element were then projected upon this phase. After this projection across all elements within the ROI, a boxcar-smoothing kernel (1.2 mm width in bin steps of 0.1 mm) was convolved with the projected amplitude data as a function of depth to obtain the laminar profile. The laminar profiles for both eye movement and visual stimulation experiments were normalized by their peak amplitudes to enable comparisons and averaging across participants.

We used bootstrapping to obtain confidence intervals on the laminar amplitude profiles in each participant and all participants combined, for both visual stimulation and eye movement experiments. For each ROI, we calculated the complex amplitudes for each run. We then resampled across runs with replacement over 5,000 iterations, and calculated the laminar profile anew for each resampled average. The 68% confidence intervals, equivalent to the standard-error-of-the-mean for normally distributed data, are shown by shading on laminar profile plots.

Peaks of the laminar profiles were calculated to quantify comparisons of depth between the attention and stimulation conditions. The peaks were also bootstrapped across the ensemble of runs to obtain confidence intervals and p-values for differences between peak values for visual stimulation and eye-movement maps.

RETINOTOPY-EYE MOVEMENT CORRELATION. Within each optimal elliptical ROI for all participants, we measured the registration between phase maps for saccadic eye movements and visual stimulation. The raw eye movement maps spanned the entire cycle (2π radians), and thus were first converted to visual field coordinates in degrees. The phase progressions within the ROI were first scaled onto a range of 90°, then these data were centered around 0° for rightward saccades, and 180° for leftward saccades. Visual stimulation phase data were corrected by estimating the hemodynamic delay as the mean phase observed in significant (*p* < 0.05) voxels of each colliculus within the elliptical ROI, and this was then subtracted from the phase data. This procedure centers the data at 0° (right horizontal), so we add 180° for the left visual field (right colliculus).

We again used bootstrapping to obtain confidence intervals on the correlations for each participant and all participants combined. For each session, we calculated a run-byrun ensemble of depth-averaged complex amplitude datasets. We then performed our correlation analysis with the retinotopy data for 5,000 averages of the attention-condition runs, each average obtained by resampling the ensemble with replacement. The *p*-values corresponded to the fraction of the correlations yielding a fit with slope ≤ 0.

### Saccadic eye movements

Eye movement were measured with the SR EyeLink 1000 Plus (Scientific Research, Ontario, Canada) both outside of the scanner during training sessions and inside the scanner during fMRI acquisition. Inside the scanner, the infrared light and camera were placed beneath the LCD display and angled at the head-coil mounted mirror allowing us to track the participant’s right eye at ~130 cm lens-to-eye distance. Raw (*x*, *y*) position coordinates were sampled at 1000 Hz. Saccade reports were generated using the EyeLink Data Viewer (Scientific Research, Ontario, Canada) and further analyzed in MATLAB. Saccades were detected using three minimum thresholds: position (> 0.15°), velocity (> 30°/s), and acceleration (> 9500°/s^2^). Eye blinks were detected when the pupil diameter was too small (< 1 mm), obstructed, or not tracked, and any saccades during blinks were discarded from analysis. Polar plots were created to represent the saccades with the direction of the saccade as the polar angle and the amplitude of the saccade as the eccentricity, which was also visualized with histograms. Smooth pursuits and correction eye movements were further distinguished from saccades by restricting saccade analysis to eye movements towards the cued horizontal visual field (i.e., left or right) and eye movements with amplitudes greater than 1° visual angle.

**Figure S1.**
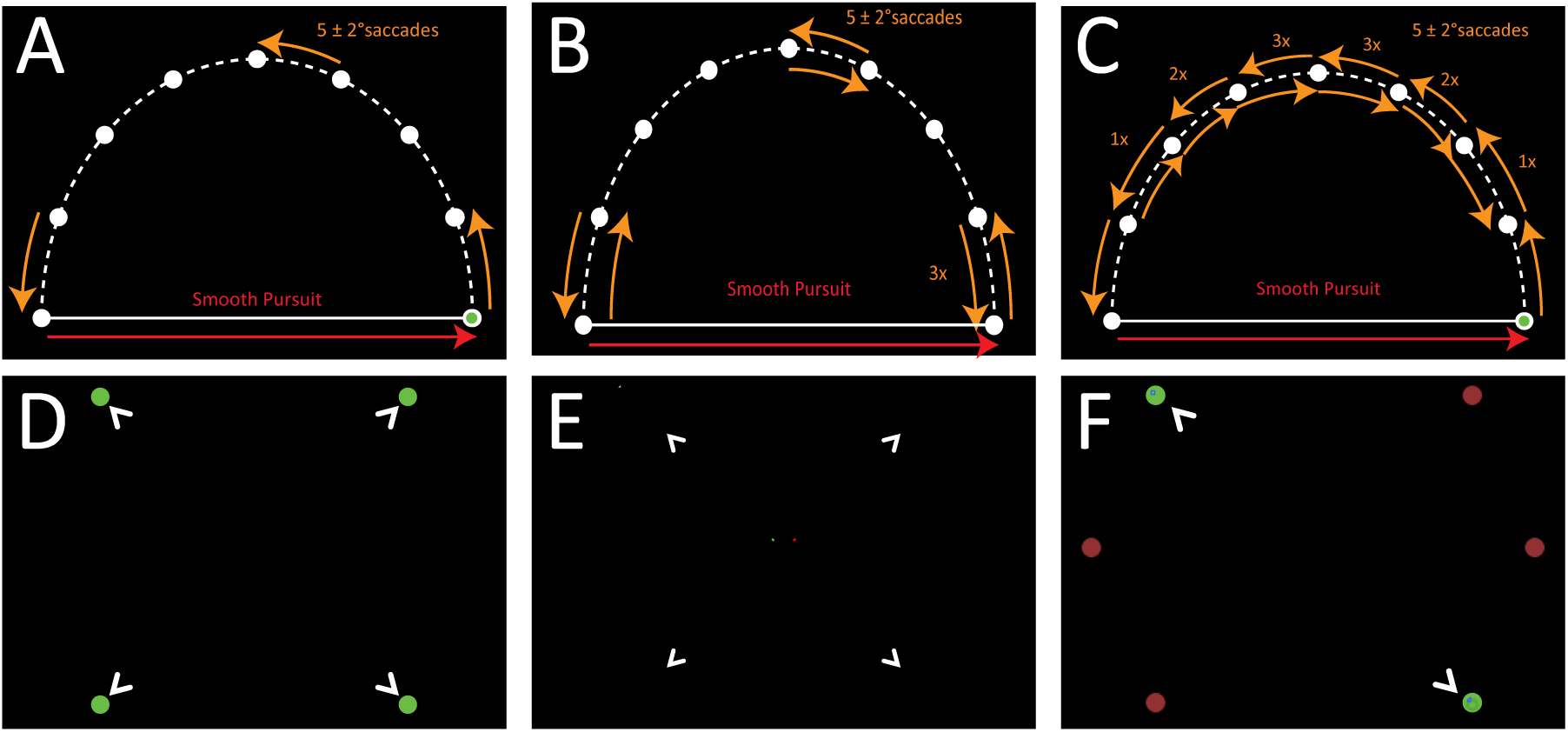
Related to Figure 1. Evolution of stimulus paradigms to evoke eye movement polar angle topography in human SC. **A.** Our first attempts to map the polar angle using a hemi-ellipse did not evoke significant activity, perhaps due to insufficient duty cycle for saccades vs. fixation. Attempts to increase the number of saccades by having participants saccade back-and-forth from previous-to-next target 7 times (**B**) or a centrally weighted number of times (**C**) also did not yield significant activity in SC, perhaps due to confounding signals from of pro- and anti-saccades occurring in sequence. We next tried to use memory-guided saccades to elicit SC activity. In one attempt, we briefly flashed a pair of peripheral green dots (**D**) for 0.2 ms and participants rapidly performed saccades between the two remembered dots for 12 s followed by saccades in the orthogonal direction. Results were still weak, but slightly improved by using foveal cues: green and red lines flashed briefly at the center of the screen to cue the direction of the memory-guided saccades (**E**). Lastly, we used a static set of red dots with a pair turned green to cue the direction of the saccades. This reduced retinal slip and increased activity significantly, and ultimately led to the design used in our main findings.

**Figure S2.**
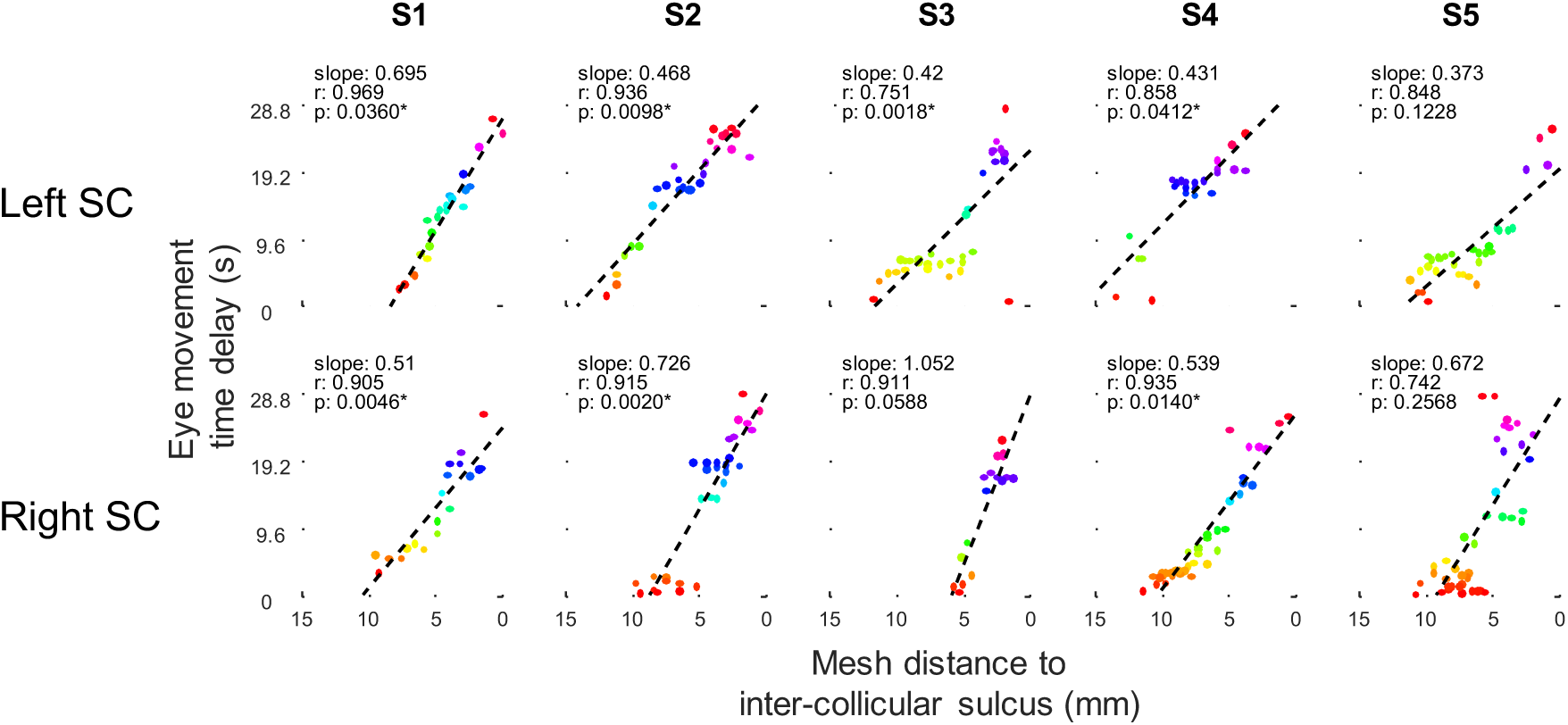
Related to Figure 6. Saccadic eye movement maps demonstrate medial-lateral progression at the individual level. The mesh distance on the surface to the inter-collicular sulcus was correlated with phase progression, represented as time delay during the saccadic eye movement task. Correlations were significant in seven out of ten sessions.

**Figure S3.**
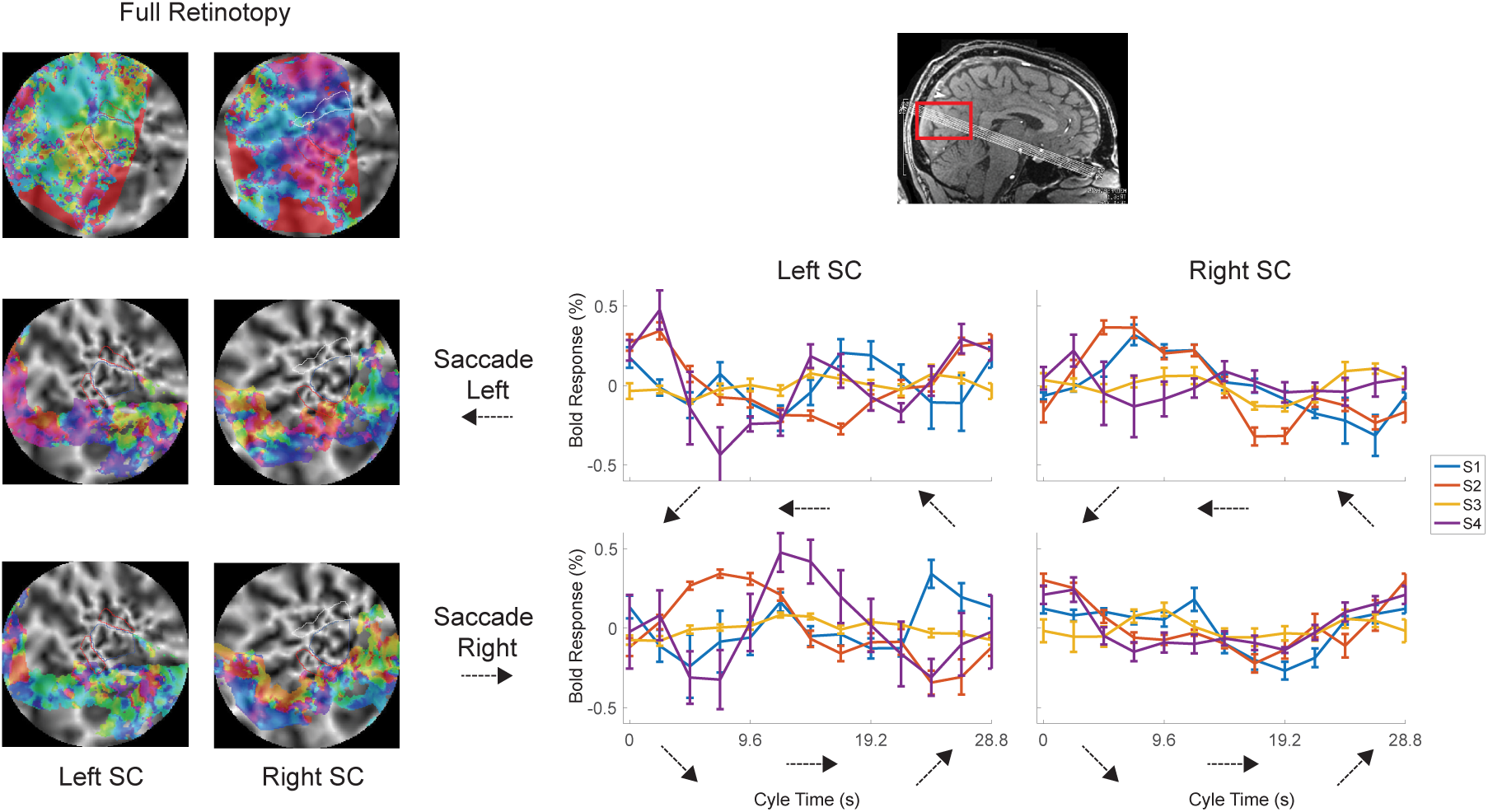
Related to Figure 5. Phase mapping of saccadic eye movements was not observed in early visual cortex. The slice prescription captured components of visual cortex (inset top right). This enabled us to explore phase progressions in early visual cortex. Four of the five participants had delineations of visual areas V1—3 as part of another experiment. A sample participant (top left) has V1 and V2 ROIs shown on the flat view. For the saccadic eye movements, we plotted phase maps on the flat view for the slices we obtained for leftward saccades (middle row) and rightward saccades (bottom row) for the same exemplar participant. We then plotted the percent modulation for the duration of the cycle (28.8 s) for all four participants with defined early visual areas. The phasic progressions do not show three distinct phases or any clear pattern across participants in early visual cortex. This analysis suggests that the phase progressions found in SC are likely not driven by visual stimulation alone.

**Figure S4.**
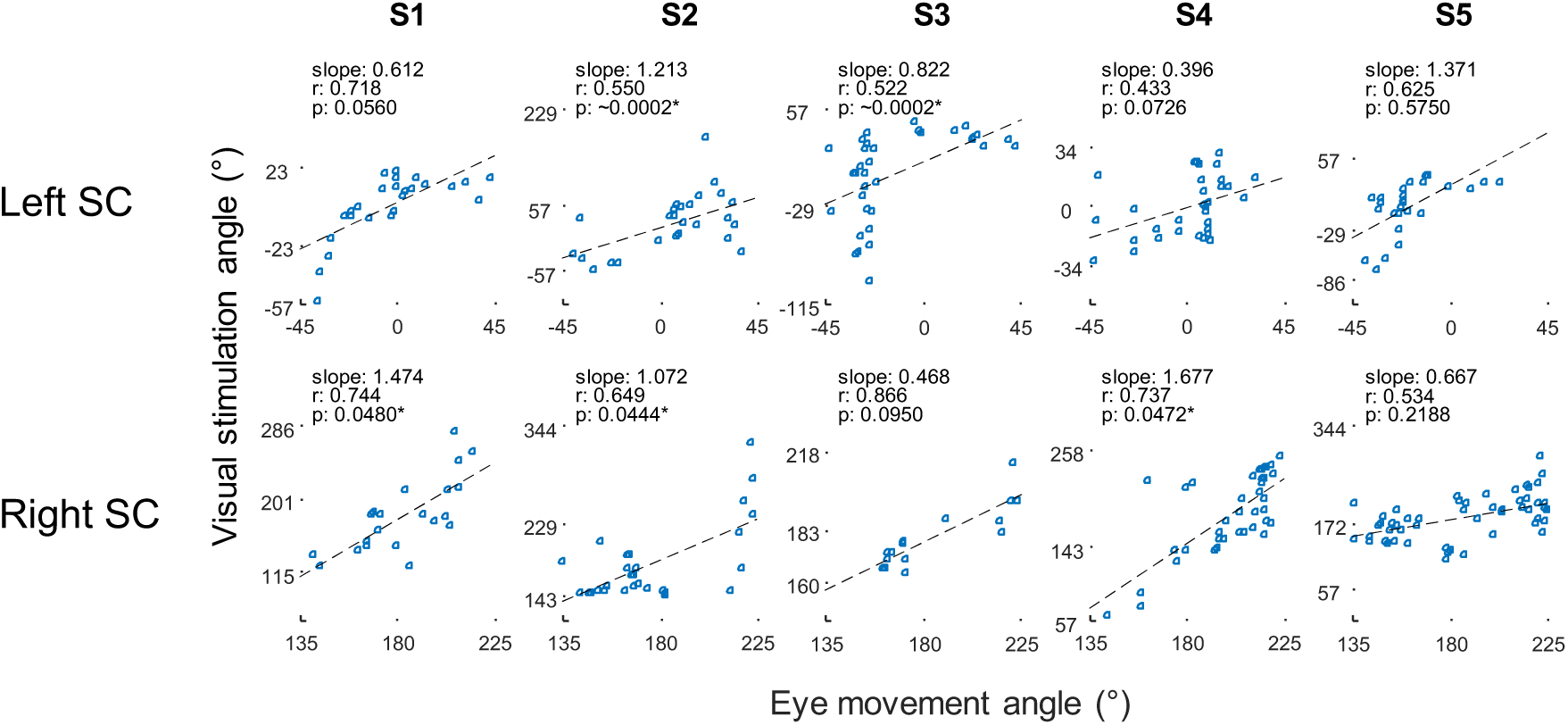
Related to Figure 7. Phase progressions for eye movement maps correlated weakly but positively with visual angle maps at the individual level. Phases from each map were converted to visual angle coordinates for comparison and correlated. P-values were obtained by bootstrapping across runs. All participants displayed a positive slope and significant in five out of 10 SCs.

**Figure S5.**
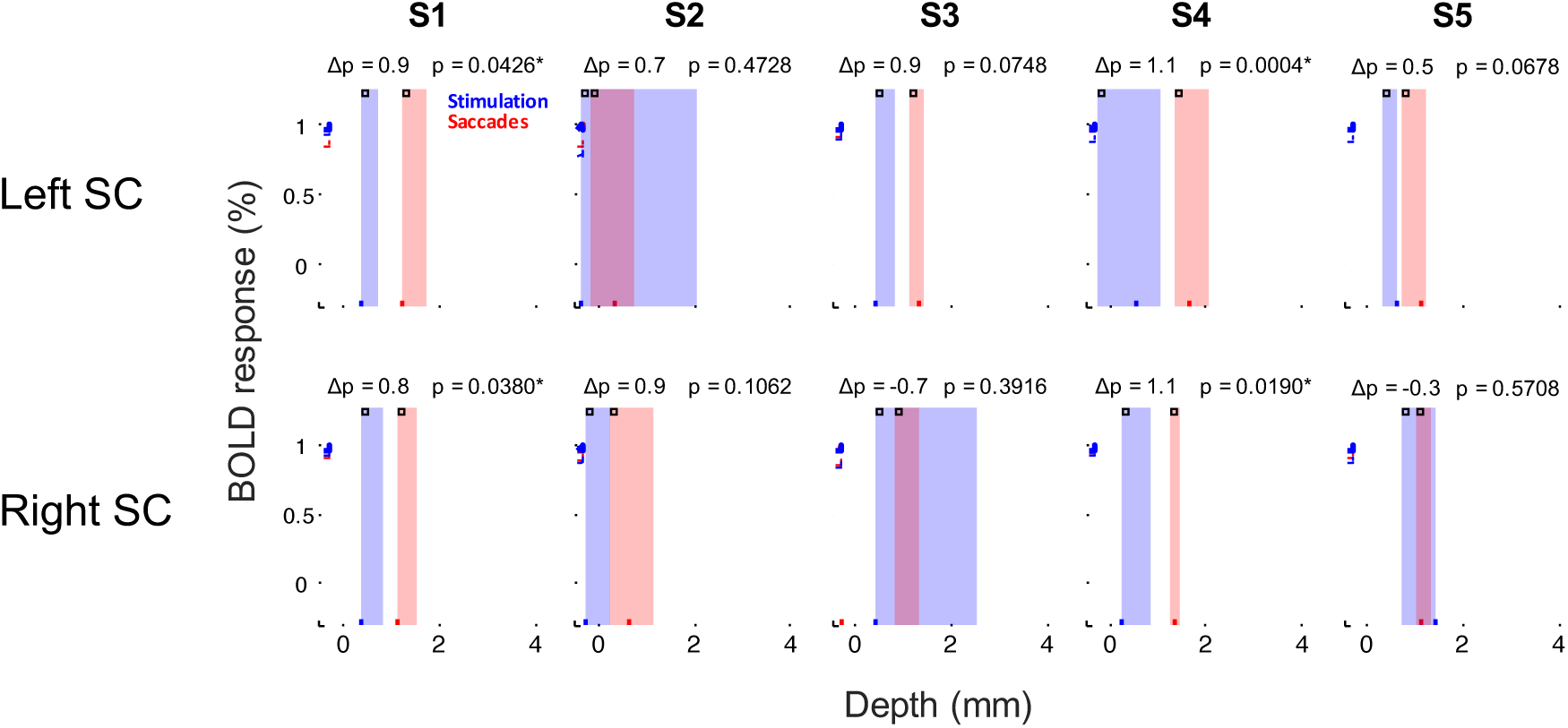
Related to Figure 8. Laminar profile depth analysis reveals activity evoked by saccadic eye movements extends significantly deeper into the SC bilaterally compared to activity evoked by visual stimulation for four out of ten sessions. Laminar profiles (solid lines) and bootstrapped confidence intervals (dashed lines) for eye movement maps (red) and visual stimulation maps (blue) are plotted. Vertical bars represent the peaks of the laminar profiles, which were significantly deeper for in four out of ten individual sessions.

## References

Adamuk, E. (1872). Uber angeborene und erworbene Association von F. C. Donders. Albrecht Von Graefes Arch. Furklinische Exp. 18, 153–164.

Apter, J.T. (1946). Eye movements following strychninization of the superior colliculus of cats. J. Neurophysiol. 9, 73–86.

Basso, M.A., and May, P.J. (2017). Circuits for Action and Cognition: A View from the Superior Colliculus. Annu. Rev. Vis. Sci.

Biotti, D., Barbieux, M., and Brassat, D. (2016). Teaching Video NeuroImages: Alternating skew deviation with abducting hypertropia following superior colliculus infarction. Neurology 86, e93–94.

Brainard, D.H. (1997). The Psychophysics Toolbox. Spat. Vis. 10, 433–436.

Büttner, U., Büttner-Ennever, J.A., and Henn, V. (1977). Vertical eye movement related unit activity in the rostral mesencephalic reticular formation of the alert monkey. Brain Res. 130, 239–252.

Chang, D.H.F., Hess, R.F., and Mullen, K.T. (2016). Color responses and their adaptation in human superior colliculus and lateral geniculate nucleus. NeuroImage.

Condy, C., Rivaud-Péchoux, S., Ostendorf, F., Ploner, C.J., and Gaymard, B. (2004). Neural substrate of antisaccades: role of subcortical structures. Neurology 63, 1571–1578.

Connolly, J.D., Vuong, Q.C., and Thiele, A. (2015). Gaze-dependent topography in human posterior parietal cortex. Cereb. Cortex N. Y. N 1991 25, 1519–1526.

Crawford, J.D., Cadera, W., and Vilis, T. (1991). Generation of torsional and vertical eye position signals by the interstitial nucleus of Cajal. Science 252, 1551–1553.

Cynader, M., and Berman, N. (1972). Receptive-field organization of monkey superior colliculus. J. Neurophysiol. 35, 187–201.

Dash, S., Nazari, S.A., Yan, X., Wang, H., and Crawford, J.D. (2016). Superior Colliculus Responses to Attended, Unattended, and Remembered Saccade Targets during Smooth Pursuit Eye Movements. Front. Syst. Neurosci. 10, 34.

Donders, F.C. (1872). Ueber angeborene und erworbene Association. Albrecht Von Graefes Arch. Für Ophthalmol. 18, 153–164.

Feldon, P., and Kruger, L. (1970). Topography of the retinal projection upon the superior colliculus of the cat. Vision Res. 10, 135–143.

Fuchs, A.F., Kaneko, C.R., and Scudder, C.A. (1985). Brainstem control of saccadic eye movements. Annu. Rev. Neurosci. 8, 307–337.

Furlan, M., Smith, A.T., and Walker, R. (2015). Activity in the human superior colliculus relating to endogenous saccade preparation and execution. J. Neurophysiol. 114, 1048–1058.

Furlan, M., Smith, A.T., and Walker, R. (2016). An fMRI Investigation of Preparatory Set in the Human Cerebral Cortex and Superior Colliculus for Pro- and Anti-Saccades. PloS One 11, e0158337.

Glover, G.H. (1999). Simple analytic spiral K-space algorithm. Magn. Reson. Med. 42, 412–415.

Glover, G.H., and Lai, S. (1998). Self-navigated spiral fMRI: interleaved versus singleshot. Magn. Reson. Med. 39, 361–368.

Hafed, Z.M., and Chen, C.-Y. (2016). Sharper, Stronger, Faster Upper Visual Field Representation in Primate Superior Colliculus. Curr. Biol. CB 26, 1647–1658.

Hanes, D.P., Smith, M.K., Optican, L.M., and Wurtz, R.H. (2005). Recovery of saccadic dysmetria following localized lesions in monkey superior colliculus. Exp. Brain Res. 160, 312–325.

Himmelbach, M., Erb, M., and Karnath, H.-O. (2007). Activation of superior colliculi in humans during visual exploration. BMC Neurosci. 8, 66.

Himmelbach, M., Linzenbold, W., and Ilg, U.J. (2013). Dissociation of reach-related and visual signals in the human superior colliculus. NeuroImage 82, 61–67.

Horn, A.K., and Büttner-Ennever, J.A. (1998). Premotor neurons for vertical eye movements in the rostral mesencephalon of monkey and human: histologic identification by parvalbumin immunostaining. J. Comp. Neurol. 392, 413–427.

Iglesias, J.E., Van Leemput, K., Bhatt, P., Casillas, C., Dutt, S., Schuff, N., Truran-Sacrey, D., Boxer, A., Fischl, B., and Alzheimer’s Disease Neuroimaging Initiative (2015). Bayesian segmentation of brainstem structures in MRI. NeuroImage 113, 184–195.

Katyal, S., and Ress, D. (2014). Endogenous attention signals evoked by threshold contrast detection in human superior colliculus. J. Neurosci. Off. J. Soc. Neurosci. 34, 892–900.

Katyal, S., Zughni, S., Greene, C., and Ress, D. (2010). Topography of covert visual attention in human superior colliculus. J. Neurophysiol. 104, 3074–3083.

Katyal, S., Greene, C.A., and Ress, D. (2012). High-resolution functional magnetic resonance imaging methods for human midbrain. J. Vis. Exp. JoVE e3746.

Khan, R., Zhang, Q., Darayan, S., Dhandapani, S., Katyal, S., Greene, C., Bajaj, C., and Ress, D. (2011). Surface-based analysis methods for high-resolution functional magnetic resonance imaging. Graph. Models 73, 313–322.

Konen, C.S., and Kastner, S. (2008). Representation of eye movements and stimulus motion in topographically organized areas of human posterior parietal cortex. J. Neurosci. Off. J. Soc. Neurosci. 28, 8361–8375.

Krebs, R.M., Woldorff, M.G., Tempelmann, C., Bodammer, N., Noesselt, T., Boehler, C.N., Scheich, H., Hopf, J.-M., Duzel, E., Heinze, H.-J., et al. (2010). High-field FMRI reveals brain activation patterns underlying saccade execution in the human superior colliculus. PloS One 5, e8691.

Linzenbold, W., and Himmelbach, M. (2012). Signals from the deep: reach-related activity in the human superior colliculus. J. Neurosci. Off. J. Soc. Neurosci. 32, 13881–13888.

Linzenbold, W., Lindig, T., and Himmelbach, M. (2011). Functional neuroimaging of the oculomotor brainstem network in humans. NeuroImage 57, 1116–1123.

May, P.J. (2006). The mammalian superior colliculus: laminar structure and connections. Prog. Brain Res. 151, 321–378.

Meredith, M.A., and Stein, B.E. (1990). The visuotopic component of the multisensory map in the deep laminae of the cat superior colliculus. J. Neurosci. 10, 3727–3742.

Mohler, C.W., and Wurtz, R.H. (1976). Organization of monkey superior colliculus: intermediate layer cells discharging before eye movements. J. Neurophysiol. 39, 722–744.

Nestares, O., and Heeger, D.J. (2000). Robust multiresolution alignment of MRI brain volumes. Magn. Reson. Med. 43, 705–715.

Qu, J., Zhou, X., Zhu, H., Cheng, G., Ashwell, K.W.S., and Lu, F. (2006). Development of the human superior colliculus and the retinocollicular projection. Exp. Eye Res. 82, 300–310.

Ress, D., Glover, G.H., Liu, J., and Wandell, B. (2007). Laminar profiles of functional activity in the human brain. NeuroImage 34, 74–84.

Robinson, D.A. (1972). Eye movements evoked by collicular stimulation in the alert monkey. Vision Res. 12, 1795–1808.

Schiller, P.H., and Stryker, M. (1972). Single-unit recording and stimulation in superior colliculus of the alert rhesus monkey. J. Neurophysiol. 35, 915–924.

Schluppeck, D., Glimcher, P., and Heeger, D.J. (2005). Topographic organization for delayed saccades in human posterior parietal cortex. J. Neurophysiol. 94, 1372–1384.

Schneider, K.A., and Kastner, S. (2005). Visual responses of the human superior colliculus: a high-resolution functional magnetic resonance imaging study. J. Neurophysiol. 94, 2491–2503.

Schneider, K.A., and Kastner, S. (2009). Effects of sustained spatial attention in the human lateral geniculate nucleus and superior colliculus. J. Neurosci. Off. J. Soc. Neurosci. 29, 1784–1795.

Sereno, A.B., Briand, K.A., Amador, S.C., and Szapiel, S.V. (2006). Disruption of reflexive attention and eye movements in an individual with a collicular lesion. J. Clin. Exp. Neuropsychol. 28, 145–166.

Sereno, M.I., Pitzalis, S., and Martinez, A. (2001). Mapping of contralateral space in retinotopic coordinates by a parietal cortical area in humans. Science 294, 1350–1354.

Singh, V., Pfeuffer, J., Zhao, T., and Ress, D. (2017). Evaluation of Spiral Acquisition Variants for Functional Imaging of Human Superior Colliculus at 3T Field Strength. Magn. Reson. Med. in press.

Sparks, D.L., and Hartwich-Young, R. (1989). The deep layers of the superior colliculus. Rev. Oculomot. Res. 3, 213–255.

Sparks, D.L., and Jay, M.F. (1986). The functional organization of the primate superior colliculus: a motor perspective. Prog. Brain Res. 64, 235–241.

Sparks, D.L., Holland, R., and Guthrie, B.L. (1976). Size and distribution of movement fields in the monkey superior colliculus. Brain Res. 113, 21–34.

Sprague, J.M., and Meikle, T.H. (1965). The Role of the Superior Colliculus in Visually Guided Behavior. Exp. Neurol. 11, 115–146.

Wade, A., Brewer, A., Rieger, J., and Wandell, B. (2002). Functional measurements of human ventral occipital cortex: retinotopy and colour. Philos. Trans. R. Soc. Lond. B. Biol. Sci. 357, 963–973.

Wandell, B.A., Chial, S., and Backus, B.T. (2000). Visualization and measurement of the cortical surface. J. Cogn. Neurosci. 12, 739–752.

de Weijer, A.D., Mandl, R.C.W., Sommer, I.E.C., Vink, M., Kahn, R.S., and Neggers, S.F.W. (2010). Human fronto-tectal and fronto-striatal-tectal pathways activate differently during anti-saccades. Front. Hum. Neurosci. 4, 41.

Wurtz, R.H., and Albano, J.E. (1980). Visual-motor function of the primate superior colliculus. Annu. Rev. Neurosci. 3, 189–226.

Wurtz, R.H., and Goldberg, M.E. (1972). Activity of superior colliculus in behaving monkey. IV. Effects of lesions on eye movements. J. Neurophysiol. 35, 587–596.

Xu, G., Pan, Q., and Bajaj, C.L. (2006). Discrete Surface Modelling Using Partial Differential Equations. Comput. Aided Geom. Des. 23, 125–145.

Yushkevich, P.A., Piven, J., Hazlett, H.C., Smith, R.G., Ho, S., Gee, J.C., and Gerig, G. (2006). User-guided 3D active contour segmentation of anatomical structures: significantly improved efficiency and reliability. NeuroImage 31, 1116–1128.

Zhang, P., Zhou, H., Wen, W., and He, S. (2015). Layer-specific response properties of the human lateral geniculate nucleus and superior colliculus. NeuroImage 111, 159–166.

